# Population imaging at subcellular resolution supports specific and local inhibition by granule cells in the olfactory bulb

**DOI:** 10.1101/046144

**Authors:** Martin Wienisch, Venkatesh N. Murthy

## Abstract

Information processing in early sensory regions is modulated by a diverse range of inhibitory interneurons. We sought to elucidate the role of olfactory bulb interneurons called granule cells (GCs) in odor processing by imaging the activity of hundreds of these cells simultaneously in mice. Odor responses in GCs were temporally diverse and spatially disperse, with some degree of non-random, modular organization. The overall sparseness of activation of GCs was highly correlated with the extent of glomerular activation by odor stimuli. Increasing concentrations of single odorants led to proportionately larger population activity, but some individual GCs had non-monotonic relation to concentration due to local inhibitory interactions. Individual dendritic segments could sometimes respond independently to odors, revealing their capacity for compartmentalized signaling *in vivo*. Collectively, the response properties of GCs point to their role in specific and local processing, rather than global operations such as response normalization proposed for other interneurons.

## Introduction

The first sensory stages of the brain are thought to format relevant environmental information in an efficient manner that facilitates easy and flexible extraction by downstream areas (Gollisch and Meister, 2010; Wilson, 2013; Wilson and Mainen, 2006). The olfactory bulb is the first synaptic station for odor information, where inputs from olfactory sensory axons are processed by a complex network of neurons before they are conveyed to higher brain areas through mitral/tufted (M/T) cells (Mori and Sakano, 2011; Murthy, 2011; Shepherd et al., 2004; Wachowiak and Shipley, 2006; Wilson, 2013; Wilson and Mainen, 2006). Interneurons in the glomerular layer may play a role in gating inputs to the olfactory bulb (Cleland, 2014; Gire and Schoppa, 2009; Shao et al., 2012; Wachowiak and Shipley, 2006), normalizing the inputs based on the overall level of input (Cleland, 2014; Olsen and Wilson, 2008) and temporal patterning of principal neuron activity (Fukunaga et al., 2012, 2014). A gain normalizing role has also been assigned to a sparse population of parvalbumin-positive interneurons in the external plexiform layer (Kato et al., 2013; Miyamichi et al., 2013).

Local bulbar processing in the deeper layers of the OB is thought to involve axon-less interneurons called granule cells (GCs), which are ~ 10 times more numerous than M/T cells (Shepherd et al., 2007). They receive synaptic inputs from M/T cells on their apical dendrites, and make reciprocal inhibitory synapses on M/T cell dendrites. These synapses mediate auto-as well as lateral inhibition across populations of M/T cells (Egger et al., 2005; Shepherd et al., 2007). The spatial extent and rules governing the connectivity between M/T cells and GCs are not well understood. In addition, since GCs can release transmitter even in the absence of action potentials (Isaacson and Strowbridge, 1998; Jahr and Nicoll, 1980; Schoppa et al., 1998), they are capable of local dendritic processing in addition to cell-wide activation (Bywalez et al., 2015; Egger et al., 2003, 2005; Zelles et al., 2006). The extent to which such local processing, which could have important consequences for the computational capabilities of the OB (Bywalez et al., 2015; McTavish et al., 2012; Wiechert et al., 2010), occurs *in vivo* remains unknown. In addition to feed-forward sensory inputs, GCs also receive extensive feedback from olfactory cortical areas (Boyd et al., 2012; Davis and Macrides, 1981; Markopoulos et al., 2012; Strowbridge, 2009).

GCs are thought to play a role in temporal patterning of activity in M/T cells (Balu et al., 2004; Dhawale et al., 2010; Egger and Urban, 2006; Fukunaga et al., 2014; McTavish et al., 2012; Schoppa, 2006), context dependent lateral interactions within the OB (Arevian et al., 2008; Cleland, 2014; Egger and Urban, 2006) and temporal patterning of mitral cell activity (Fukunaga et al., 2014; Lagier et al., 2007). At a behavioral level, the extent of their activation can influence odor discrimination times (Abraham et al., 2010; Nunes and Kuner, 2015). Theoretical and computational studies have also suggested a role for GCs in pattern separation and decorrelation (Gilra and Bhalla, 2015; Wiechert et al., 2010). There is, however, some uncertainty about the extent of GC influence on some aspects of MC activity (Fukunaga et al., 2014). In this regard, cortical feedback is thought to flexibly alter the activity of GCs based on experience and context (Boyd et al., 2012; Kato et al., 2012; Markopoulos et al., 2012; Otazu et al., 2015; Restrepo et al., 2009). GC activity *in vivo* has been recorded using electrophysiological techniques, but the relatively low yield of this technique has prevented a comprehensive view of the repertoire of GC responses to odors (Cang and Isaacson, 2003; Cazakoff et al., 2014; Labarrera et al., 2013; Tan et al., 2010; Wellis and Scott, 1990). Recent experiments using multiphoton imaging of populations of GCs have revealed interesting features of GC activity, for example that they are weakly and sparsely activated in anesthetized mice, and become more densely activated in awake mice (Kato et al., 2012), a finding confirmed by electrophysiological recordings (Cazakoff et al., 2014).

Despite the welcome resurgence of research on GC function, none of these studies have shed light on how GC activity relates to the statistics of inputs to the OB (glomerular activation patterns, for example), nor have they examined the subcellular local activation of GCs in vivo. More generally, many key questions about the spatiotemporal dynamics of activity in GCs and their relation to sensory stimuli remain to be addressed. Here, we have used multiphoton microscopy and calcium imaging to examine odor responses in a large population of GCs, both in their somata as well as in their dendrites.

## Results

We used viral expression of GCaMP3 (Tian et al., 2009) to investigate odor-evoked activity in GCs of adult mice. In a few experiments that are flagged clearly, we also used a more sensitive indicator GCaMP5 (Akerboom et al 2012). To assist in identification of infected cells, we also co-expressed dTomato in these cells using the 2A viral element (Figure 1). We injected AAV2/9 virus into one or both olfactory bulbs, and waited for 3 weeks or more before imaging. Infected OBs were visualized after removal of the overlying bone with wide-field fluorescence imaging (Figure 1A).

**Figure 1:**
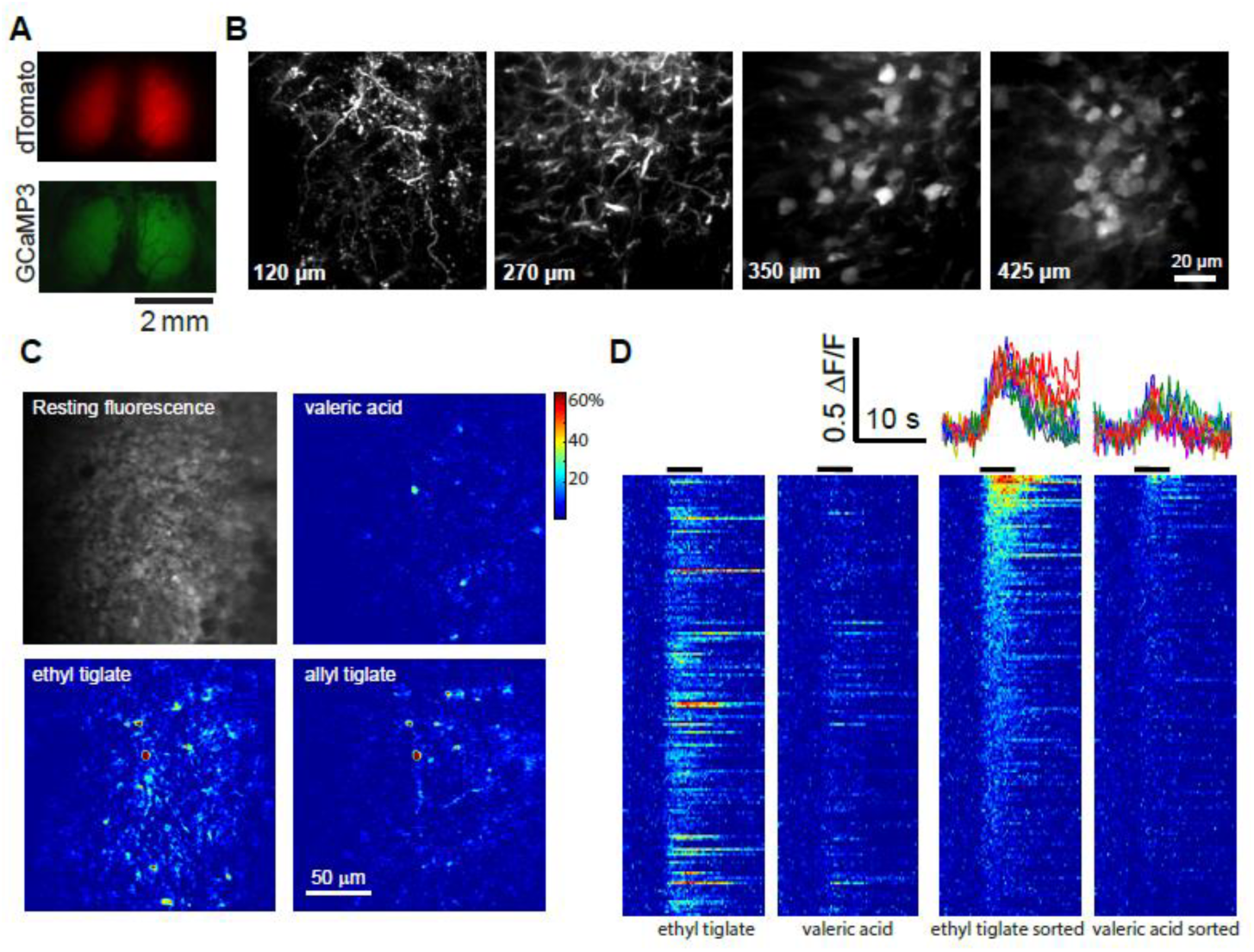
Imaging odor responses in GCs. A. OBs from living mice imaged through a cranial window after viral expression of dTomato and GCaMP3 (top). B. Labeled GCs imaged (dTomato channel) in vivo using two-photon microscopy. Optical sections at different depths reveal dendrites, spines and cell bodies. Expression is sparse in this example because of low volume of virus injection. C. Odor-evoked responses in cell bodies shown as differential fluorescence images. D. Odor-evoked responses to 2 odors in 169 cells from a typical experiment. For each odor, responses are shown in arbitrary order in the left panels and in descending order of response amplitudes on right. Traces for 5 selected GCs are shown top right for 2 odors. Odor stimulus duration indicated by black lines above colormap panels.

Immunohistochemical analysis revealed that GCs can be readily labeled with dTomato and GCaMP3, with negligible infection in mitral cells, at least with AAV2/9 at the titers used in this study (Figure 1A, Supplementary Figure 1). Even if a few mitral cells were labeled, GCs could be readily identified by their smaller size and deeper locations.

We used multiphoton microscopy to image GC somata and apical dendrites in anesthetized mice. In the external plexiform layer, dendrites and spines could be readily identified in the red channel (Figure 1B), and somata of GCs could be discerned up to 400 μm below the pial surface (Figure 1C; Supplementary Movie 1). Although only a few GC somata were labeled in the example in Figure 1B, GCs could be labeled at varying densities by titrating the amount of virus – for experiments on somatic imaging we used high titers for dense labeling. Odors were delivered using a custom-built olfactometer and GCs were imaged at a sampling rate of 4-8 Hz (see Methods). Odor stimulation caused an increase in fluorescence in GC somata (Supplementary Movie 2), with the number of responding cells varying with odor identity (Figure 1C). In addition to discrete somatic regions, neuropil responses could often be observed, presumably due to dendrites of GCs located in deeper layers.

Somatic regions were chosen for analysis based on the bright fluorescence in the red channel, and fluorescence changes in the GCaMP channel monitored. Responses in individual GCs were reliable (Supplementary Figure 2A) and those indifferent GC somata had differing amplitude and time course (Figure 1D), which were dependent on the odorant. For each odorant, we ordered the responses by the average amplitude to visualize the extent of activation more clearly (Figure 1D, right panels). This method of visualization demonstrates the diversity of temporal dynamics, which does not just depend on the response amplitude. For example, responses that are sustained beyond the odor presentation occur for different amplitudes. We use this mode of representation in subsequent figures for visual clarity.

**Figure 2:**
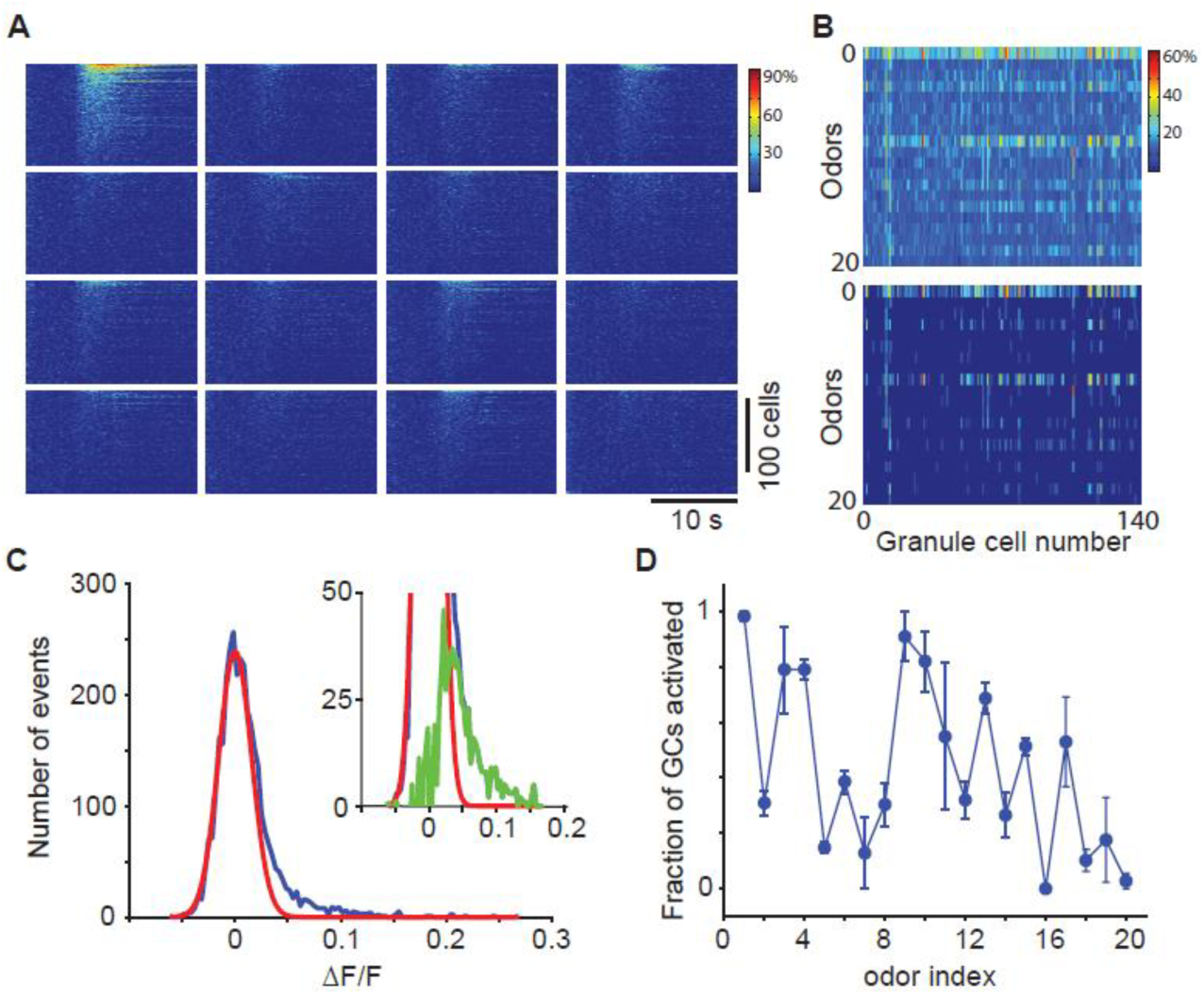
Characterizing GC responses. A. Responses of 169 GCs to 20 odors from the experiment shown in Figure 1D. For each odor, GCs are arranged in descending order of their response amplitudes, therefore the order across odors could be very different. B. Matrix of average responses of the 169 GCs to 20 odors. Top panel shows all the average Δ F/F values and the bottom matrix shows only those responses that are above a threshold of 0.05. C. Distribution of average Δ F/F values for 3380 GC-odor pairs (blue) and the noise distribution calculated from baseline measurements (red). nset shows the same distributions on an expanded scale, with the signal distribution (green) calculated as the difference between the blue and red distributions. D. The fraction of GCs in imaged regions that were activated each of the 20 odors, normalized to that for odor 1. Error bars are standard deviations from 3 experiments.

### Characterizing odor-evoked responses

We used a suite of 20 odorants to examine the responses of the GC population. The odorants were chosen to activate the dorsal surface of the mouse OB at differing densities (see below). Figure 2A shows that different odors activated different fraction of GCs in the imaged area. A matrix of average response amplitudes illustrates this clearly (Figure 2B), with some odors (e.g., ethyl tiglate) activating many GCs in the imaged region, while other odors activating hardly any cells. Visualizing only responses that were above a hard threshold value (0.05 in this example) makes the odor-dependent differences in the number of responding GCs clearer (Figure 2B).

While stronger responses could clearly be distinguished from noise, weaker responses were harder to discern. To obtain an objective measure of the distribution of responses, we plotted the histogram of responses of all GC-odor pairs in a given experiment. We used the baseline fluorescence period to obtain a noise distribution for the DF/F measurements and fitted a normal distribution to it (Figure 2C, red line). Deviations from this normal distribution are due to odor-evoked responses (Figure 2C, green line in the inset). The signal and noise distributions are highly overlapping, and a significant portion of responses will be lost using a hard threshold (for example, 2 SD of the noise distribution). Although this cannot be avoided for identifying individual responses, we can estimate the total number of responses for each stimulus by subtracting out the noise distribution. Such a population analysis indicated that the fraction of GCs in the imaged region that are activated varies widely for different odors (Figure 2D).

We performed several control experiments to rule out some biases in our recordings. First, we found the responses of individual GCs across multiple trials to be reliable (Supplementary Figure 2A). Second, we found no systematic relation between the resting fluorescence of a GC and the maximal ΔF/F (Supplementary Figure 2B), ruling out a dominant effect of expression level of GCaMP3 on responses. Third, we found that most of the imaged GCs are capable of responding to stimuli once GABAergic inhibition is attenuated pharmacologically (Supplementary Figure 2C, Supplementary Movie 3), indicating that the lack of activity in a majority of GCs is due to insufficient synaptic drive. Finally, we also performed additional experiments with a newer generation indicator GCaMP5 (Supplementary Figure 3), as well as other experiments in awake animals (Supplementary Figure 4) which confirmed the basic conclusions of the study.

**Figure 3:**
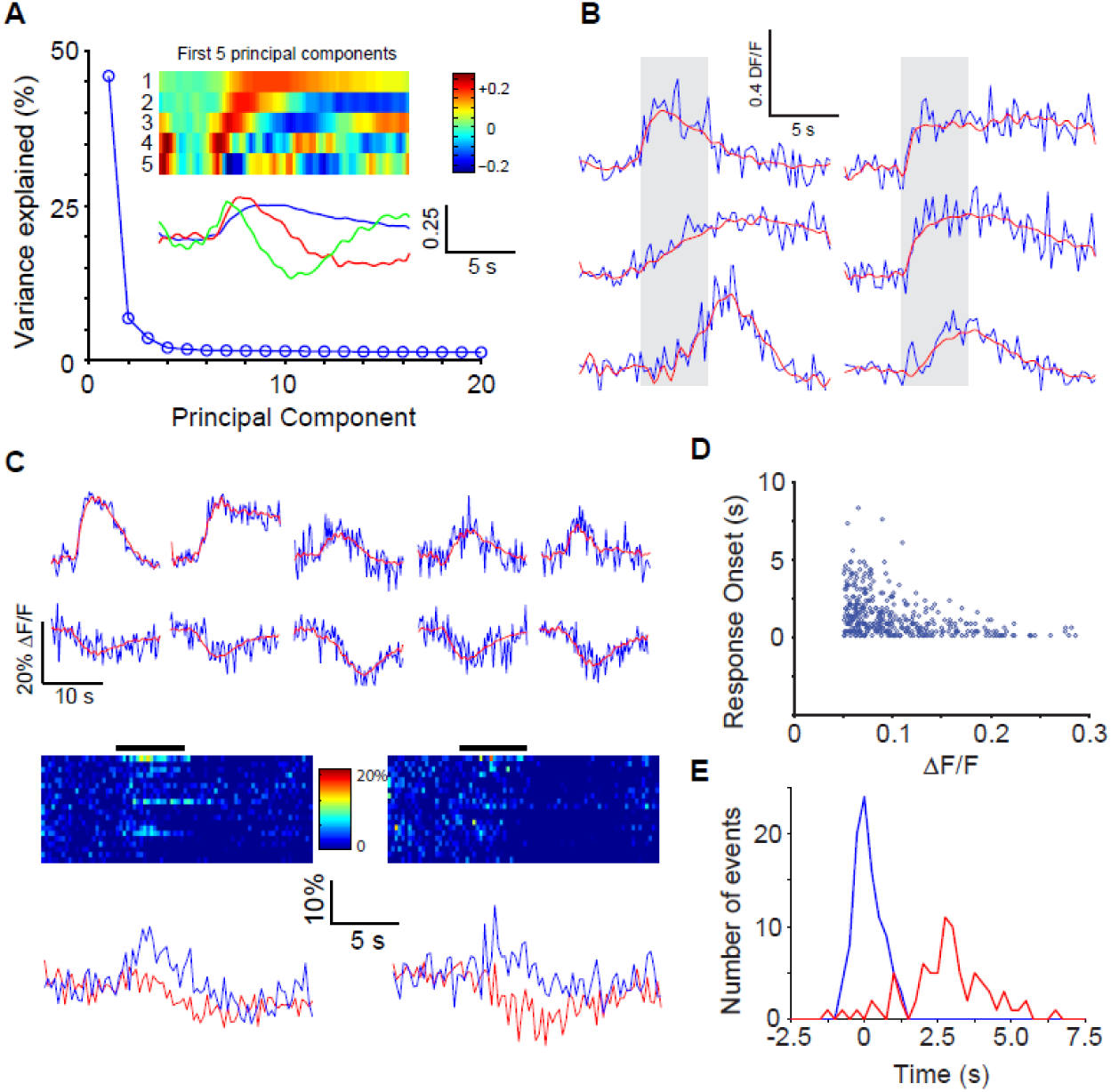
Temporal dynamics of GC responses. A. PCA of the temporal profiles of responses of all GCs to all odors revealed that the first few principal components could explain most of the variance in the responses. The first 5 components are shown as color maps and only 3 components are shown as line traces below for clarity. B. Examples of the time course of a few responses in GCs, with fits from the first 5 PCs (red) superimposed on raw fluorescence traces (blue). C. Examples of inhibitory responses, along with additional examples of excitatory responses in the same experiment. Inhibitory responses could be well-explained by the same 5 PCs used for excitatory responses. The responses of two exemplar GCs to all 20 odors reveal that the same GC can have both excitatory and inhibitory responses to different odors. Selected traces are shown below for clarity. D. Response latency (see Methods for details) for excitatory responses plotted against the average response amplitude (top) shows that strong responses had less variable, short latencies. E. The distribution of response latency for inhibitory responses (red) was significantly (p < 0.05) shifted to the right compared to excitatory responses (blue).

**Figure 4:**
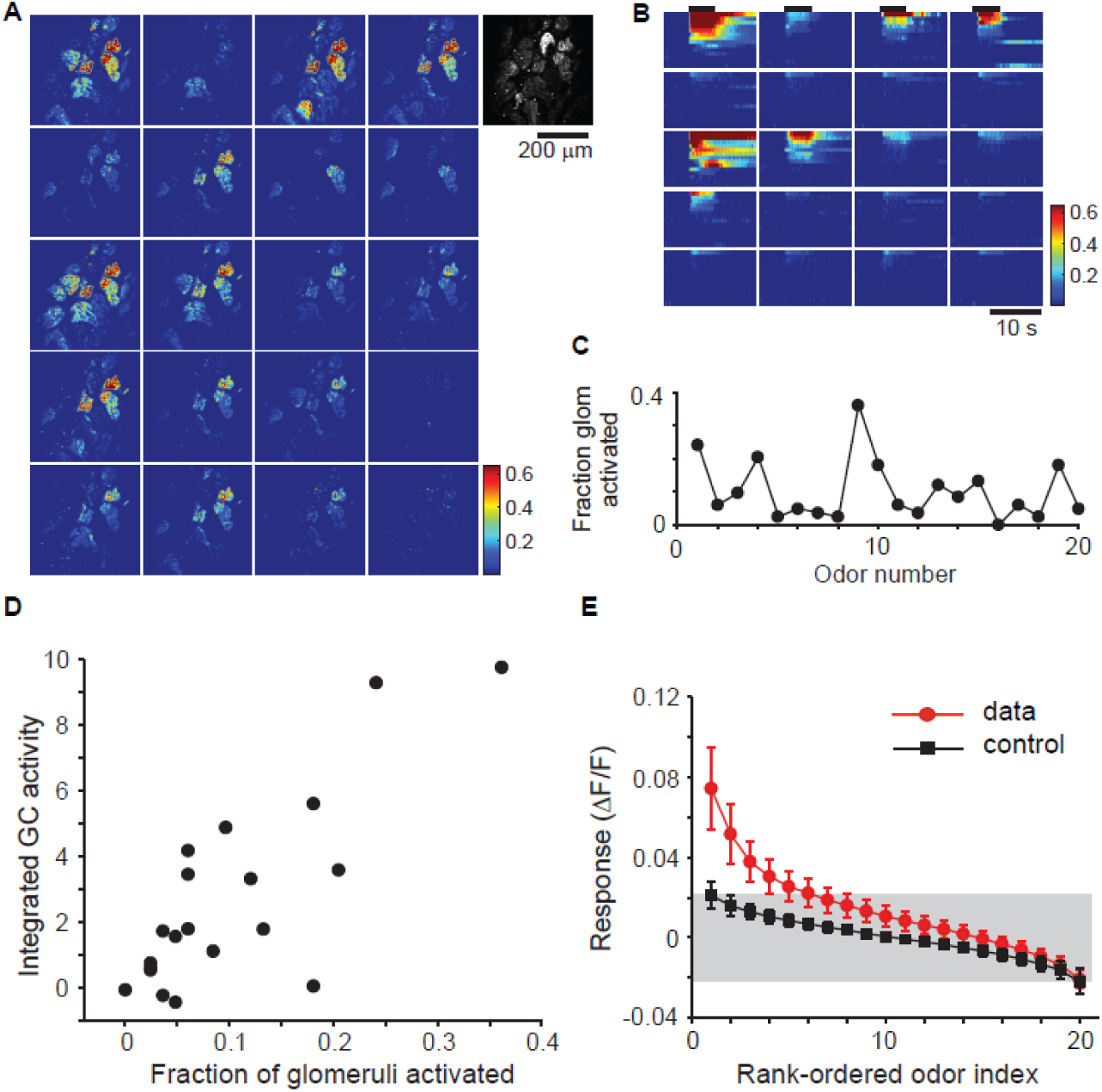
Response sparseness correlates with glomerular activation pattern. A. The extent of glomerular activation by 20 odors was estimated by imaging responses in OMP-GCaMP3 mice. Fractional fluorescence changes for each of the odors are shown for one experiment, with the resting fluorescence shown at right in grayscale. B. Time course of responses of individual glomeruli to all 20 odors (top). C. Fraction of glomeruli activated (see Methods) varies widely for different odors. D. The integrated activity of a population of GCs for a given odor is highly correlated with the fraction of glomeruli activated by that odor (representative experiment, averages in text). E. Rank-ordered odor responses of GCs reveal that only a few odorants activate any given cell strongly. For each GC, the absolute values of responses were ranked in descending order and then the average across all GCs was obtained. Only the first 3 rank-ordered responses are significantly (p < 0.05) different from control plots from simulated data derived from the experimentally-observed noise distribution.

### Temporal structure of responses

Although electrophysiological recordings have excellent temporal resolution, they are technically challenging and offer sparse sampling. We leveraged the high numbers afforded by population imaging to provide a comprehensive view of GC dynamics.

The example recordings shown above (Figures 1 and 2) hint at the diversity in the dynamics of odor responses. These included transient, accumulating, steady and delayed responses. To obtain an unbiased measure of the types of different temporal dynamics, we performed principal component analysis (PCA) of all responses (all GC-odor pairs) within a given experiment. We found that the first few principal components explained a large fraction of the variance in the time course of responses (Figure 3A). A diversity of responses could be described well using 5 PCs (Figure 3B). In addition to increases in fluorescence intensity upon odor stimulation, we also noticed reductions in fluorescence in some cases (Figure 3C). The response dynamics of a given neuron to different odors could be very different, with both inhibitory and excitatory responses (Figure 3C, bottom). Inhibitory responses were far less frequent than excitatory responses (1061/8000 cell-odor pairs or 0.133 ± 0.008 excitatory responses vs. 83/8000 cell-odor pairs or 0.010 ± 0.002 inhibitory responses; p < 0.001), but the proportion of inhibitory responses will be an underestimate because of the low sensitivity of calcium imaging to hyperpolarization. The GABAergic antagonist bicuculline applied to the surface of the OB (see Methods) greatly reduced the fraction of GC responses that showed odor-evoked reductions (9/2888 or 0.003 ± 0.001 of cell-odor pairs, p < 0.001, Fisher’s exact test, compared to the fraction without bicuculline above), indicating that the inhibitory responses are largely due to ionotropic GABAergic mechanisms.

We noticed that several odor responses occurred with significant delays following odor presentation (Figure 3B). These delays were not due to stimulus dynamics because for the same stimulus, different GCs in the same imaged area could respond with different delays. When the delay was plotted as a function of response amplitude, it became clear that delayed responses were much more prevalent when the magnitude of the response was smaller (Figure 3D). Such a relation could arise by either difficulty in detecting responses when the amplitude is small due to poorer signal-to-noise ratio. Alternately, there could be a threshold effect whereby smaller amplitude responses are typically a result of slow accumulation of synaptic input. Our data rule out the possibility that strong synchronous inputs arrive in a delayed manner after a period of quiescence. Instead, it would seem that if a GC receives strong excitation, it does so early on. We also found that inhibitory responses occurred, on average, with a greater delay than excitatory responses (Figure 3E). The average delays for the strongest 100 positive and 80 negative responses were 0.13 ± 0.48 and 2.78 ± 1.43 seconds respectively (p < 0.001; t-test).

Our data indicate that odor responses in GCs are heterogeneous and can be explained by a small number of elementary components. Interestingly, our imaging method is sensitive enough to detect inhibitory responses, which typically arise at longer delays following stimulation.

### Density of GC activation depends on stimulus density

Different odorants activate differing number of GCs (Figure 2). We wondered if there was a relation between the density of GC activation and the degree of activation of inputs to the OB. We first characterized the spatial extent of glomerular activation by the panel of odorants used in this study using mice expressing GCaMP3 in OSNs (Isogai et al., 2011; Rokni et al., 2014).

We imaged the glomerular inputs in OMP-GCaMP3 mice and recorded responses over a large region of the dorsal surface. Robust odor-evoked responses were imaged in the glomerular layer of OMP-GCaMP3 mice to many of the odors (Figure 4A). The time course of responses varied depending on the glomerular and odor identities (Figure 4B). By identifying all glomeruli in the imaged region using the resting fluorescence of GCaMP3, we estimated the fraction of glomeruli activated by a given odor. Averaging across multiple regions of interest and animals, we determined that odorants in our panel activated between 0 and 30% of glomeruli imaged on the dorsal surface (Figure 4B,C).

We next asked whether the density of responses of GCs was related to the fraction of activated glomeruli. Such a relation is inevitable in a purely feed-forward network, but lateral interactions within the OB could lead to winner-take-all-like activation, where a similar number of GCs could be activated by different odorants (Koulakov and Rinberg, 2011). We used two measures to represent the density of activity of GCs. First, we simply summed the responses of all labeled GCs to each odor, and plotted this summed activity against the fraction of glomeruli activation (Figure 4D). The summed GC activity was strongly correlated with glomerular activation (correlation coefficient of 0.80 in example shown in Figure 4D), with an average correlation coefficient of 0.80 ± 0.05 (SD, N = 7 experiments). A similar strong correlation was also found when considering the fraction of active GCs in the imaged field of view (0.81 ± 0.04, N= 7 experiments). These strong correlations were also present if we used a modified metric for glomerular activation that took into account the amplitude of glomerular responses, rather than treating them in an all-or-none manner.

A second measure of GC activation we used is the population sparseness, which is related to the fraction of GCs that are activated strongly by a given stimulus (Rolls and Tovee, 1995; Willmore and Tolhurst, 2001). There are multiple ways of characterizing population sparseness (Willmore and Tolhurst, 2001), and we chose a metric that is designed for responses involving only positive values (Rolls and Tovee, 1995; Willmore and Tolhurst, 2001)(see Methods). We found that this measure was also highly correlated with the glomerular input activity (Supplementary Figure 5), with an average correlation of 0.78 ± 0.04 (N = 7 experiments). These data indicate strongly that the extent of activation of the GC population is determined in large part by the density of sensory input.

**Figure 5:**
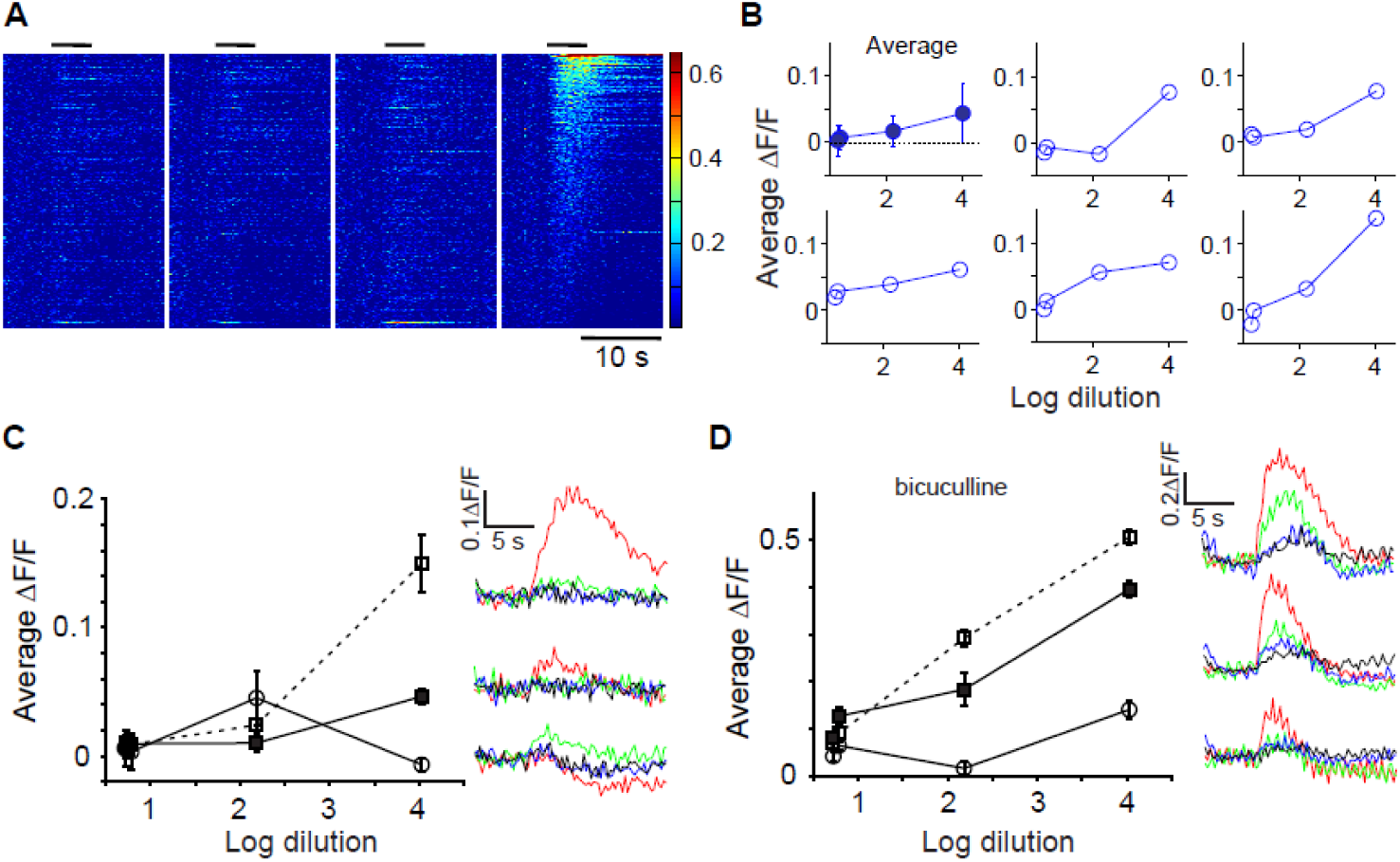
. Effects of increasing concentration on GC responses. A. Responses of all GCs from one experiment to four concentrations of allyl tiglate. Responses were arranged in descending order of amplitude for the highest concentrations and the rest of the plots followed the same GC rank order. B. The average response to the four concentrations across the entire population (top left) and 5 randomly picked individual GCs. C. Based on the highest concentration, 3 groups of 5 GCs were picked as the highest-amplitude responders, medium responders and the lowest amplitude responders. The responses of these three groups of GCs were then averaged for all concentrations, revealing non-monotonic responses as concentration increased. Time course of responses to the 4 concentrations are shown for the 3 groups at right (black, blue, green & red colors for increasing concentration). D. A similar analysis in a different experiment performed after bicuculline was added to the surface of the OB showed no evidence of non-monotonic responses.

We next asked how responsive individual GCs are to odor stimuli. We arranged the odor tuning curve for each GC in a descending order of the response amplitude, and averaged these across all GCs in an experiment (Figure 4E). Because of the strict rank ordering, the tail of this curve reflects a biased noise distribution, which can be estimated using Monte Carlo sampling from the measured noise distribution (see Methods). Comparing the data distribution with the null distribution revealed that on average a GC responded to 3 odors out of the 20 used (Figure 4E). We also calculated a widely-used measure called lifetime sparseness (Willmore and Tolhurst, 2001), which does not rely on thresholds to define responses. This measure varied widely from cell to cell (Supplementary Figure 5), but on average this measure confirmed the sparseness of GC responses noted above.

The measure of sparseness of GC responses is likely to be affected by the sensitivity of the reporter used. We performed additional experiments using a more sensitive indicator, GCaMP5 (Supplementary Figure 5). Although more GCs were indeed activated by odors, the additional data confirmed the basic conclusion that the density of responses is strongly correlated with the extent of glomerular input (Supplementary Figure 5).

These results indicate that the density of GC activation is related strongly to the extent of glomerular activation, and GCs can be readily recruited even in anesthetized animals through dense activation of glomeruli.

### Concentration dependence of GC activity

We found that there was a strong correlation between the extent of glomerular activation and the density of active GCs. We wondered if this relation also applies to increasing concentration of a given odor, a more straightforward way of increasing stimulus strength than altering odor identity to activate more glomeruli (Figure 5).

Our stimulus set included 4 concentrations of allyl tiglate, spanning 3 orders of magnitude as measured by a miniPID device. We first confirmed that glomerular input responses to these four concentrations were strictly monotonically increasing (Supplementary Figure 6). We then sorted GC responses to the highest concentration of odorant and displayed the responses to other concentrations in the same order (Figure 5A). Visually, it becomes clear that responses are stronger and more GCs are activated by increasing concentration. This can be quantified by taking the average fluorescence change across all GCs (Figure 5B, upper left panel), revealing a monotonic relation. Individual GCs display a range of dependence on concentration (Figure 5B). We wondered whether all GCs have monotonic relation to odor concentration. To examine this issue objectively, we pooled GCs at the extreme of responses to the strongest concentration: 5 GCs that had the strongest responses, 5 GCs with responses in the middle range and 5 GCs with the weakest responses (or the most inhibited ones to be precise). We then examined the dependence of concentration of the responses of these GCs. This analysis uncovered a clear non-monotonic relation in a subset of GCs (Figure 5C), with the middle concentration giving rise to larger responses than the lower or higher concentration.

Since glomerular activation, at least in our experiments, always increased monotonically or saturated with odorant concentration (Supplementary Figure 6), we reasoned that non-monotonic changes in individual GC activation are likely to be due to circuit interactions in the OB. In particular, it is possible that increasing activation of more glomeruli with increasing concentration could lead to lateral interactions within the OB that involve inhibition. We examined the concentration-dependence of GC responses in experiments where bicuculline was infused over the OB. We found that the non-monotonic relation observed for some GCs in control conditions was no longer found in experiments where inhibition was blocked with bicuculline (Figure 5D).

Our data indicate that although many GCs are proportionally recruited by increasing stimulus intensity, other GCs can be inhibited at higher activation levels, leading to non-monotonic responses.

### Spatial organization of responding GCs

We exploited our ability to record hundreds of GCs at once, to ask two questions about their spatial organization. First, are GCs that are close together functionally more similar than those located farther away? Second, are GCs responding to activation of a single glomerulus located in close proximity?

To answer the first question, we adapted an analysis method developed previously by us (Soucy et al., 2009). In this method, we characterized the functional similarity of pairs of GCs by the cross correlation of their odor response spectra. GCs that respond in the same way to a panel of odors will have a correlation value close to 1, those that respond in an uncorrelated manner will have a value close to 0 and those that are anti-correlated will have a value of ‐1. We then plotted this similarity value for pairs of glomeruli as a function of their physical separation (Figure 6A). Analysis over a large number of pairs revealed no trend – that is, GC pairs separated by almost 300 microns were just as similar as those separated by a few microns (Figure 6B). We compared this relation to a control one in which we preserved the distribution of spatial location of responding cells, but scrambled their response spectra. The observed relation was very similar to the control distribution, but had a slight dependence on distance (Figure 6B).

Since neighboring glomeruli can be functionally quite dissimilar (Bozza et al., 2004; Ma et al., 2012; Soucy et al., 2009), any spatial clustering of responding GCs could be obscured by overlapping populations of GCs activated by neighboring glomeruli. We wondered if sparse activation of glomeruli may reveal clustered activation of GCs. Visual examination of the locations of active GCs suggested that odors with sparse glomerular activation (as characterized above in Figure 3) activate a small number of GCs that appeared to be clustered within the imaged region (Figure 6C). To quantify this impression, we calculated the average pair-wise separation between activated GCs for each odor and examined its relation to the sparseness of glomerular activation (Figure 6D). For odors that activated glomeruli sparsely, the average pairwise distance reached an asymptote of ~127 μm, which was the expected value for random pairs of GCs selected from the imaged region (dashed line in Figure 6D). Interestingly, the pairwise separation for GCs activated by “sparse” odors was smaller than those for denser odors (p < 0.001 with one-way ANOVA; p < 2e^-6^ for a linear correlation analysis). This drop in pairwise separation was not an artifact of small numbers, since simulations of sparse random sampling from the population of imaged GCs led to higher variability in individual experiments, but not smaller separation (Supplementary Figure 7).

**Figure 6:**
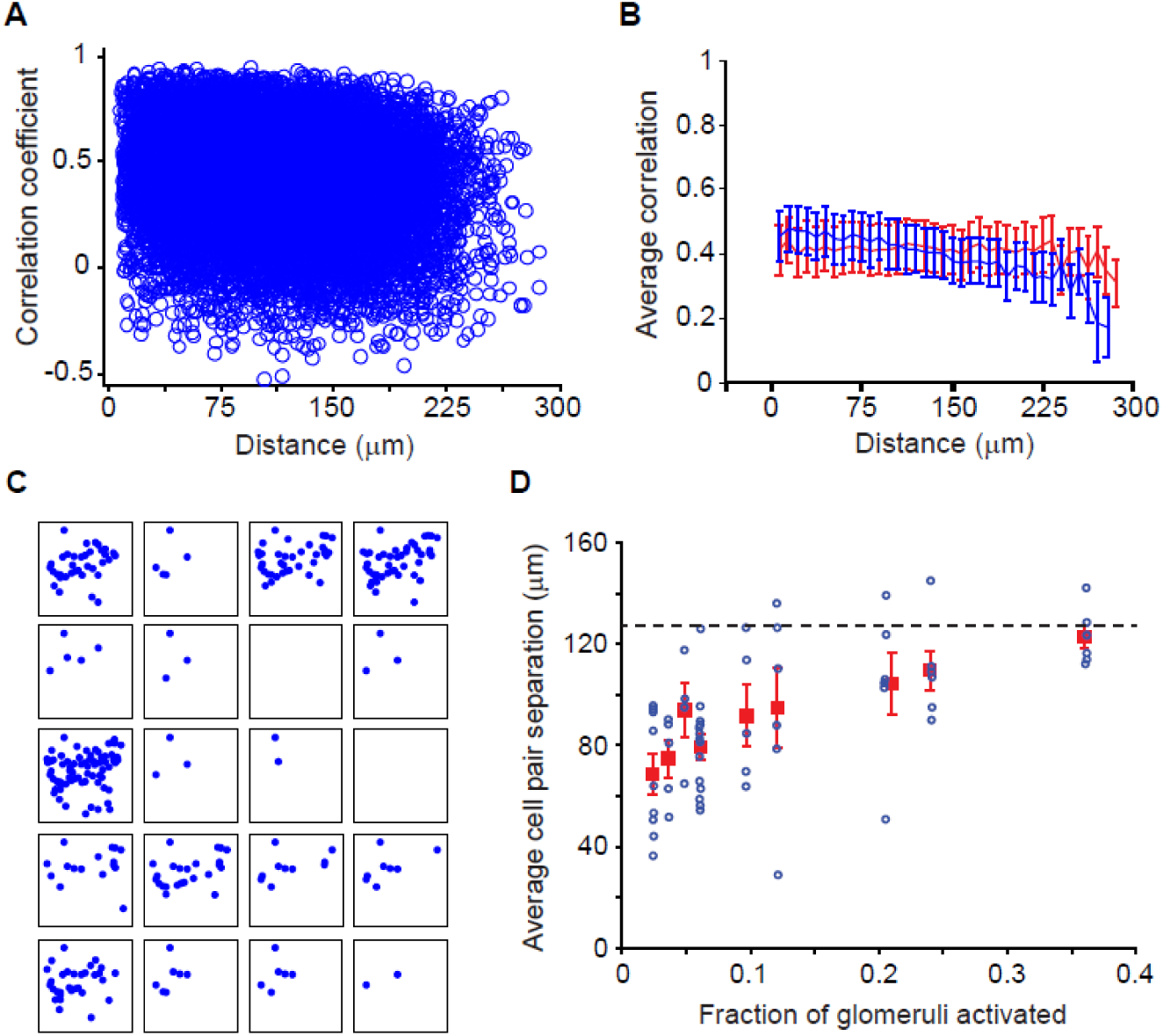
Spatial organization of responding GCs. A. Scatter plot of similarity of GC odor tuning between a pair of GCs (calculated as cross correlation of their odor responses - see methods) and their spatial separation across all experiments. B. The average similarity varies very little with separation, and is not significantly different from a control values calculated by randomizing odor response across GCs (red). C. The spatial locations of active GCs for each of the odors in one experiment. D. The average pair-wise separation of GCs was calculated for each odor and plotted against the glomerular activation fraction for that odor. Blue circles are from individual experiments and averages are in red squares. Errorbars are SEM. The dashed line is the expected average separation if GCs are activated randomly in terms of their spatial location. The trend for lower separations for sparsely-activating odors was significant (p < 0.01).

These data suggest that GCs activated by a glomerulus in isolation are clustered, which could arise from either selective connectivity or from simple geometric considerations (Murthy, 2011) (see Discussion).

### Dendritic responses

Since GCs release neurotransmitter from their dendrites, their output can be modulated by local depolarization as well as cell-wide depolarization. Experiments in slices have indicated that both modes of depolarization can exist (Bywalez et al., 2015; Egger et al., 2005), but the occurrence of dendritic depolarization in vivo has remained in question. Imaging allowed us to examine directly how the apical dendrites of GCs respond to odorants.

We chose to first examine GC dendrites as a population by densely labeling GCs selectively using the VGAT-Cre mouse line (Vong et al., 2011). By injecting virus to conditionally express GCaMP3 only in GABAergic interneurons, we ensured that there was no contribution of M/T cell dendrites when imaging was done in the EPL. Robust odor-evoked responses were observed in the EPL (Figure 7A,B). The amplitude of population responses in the EPL was strongly correlated with the density of glomerular activation (Figure 7C; correlation coefficient 0.79 ± 0.08), just as it was for GC somata. In some experiments, we examined responses in GC somata as well as in the EPL in the same preparation. The integrated activity was highly correlated in the two regions (Figure 7D).

**Figure 7:**
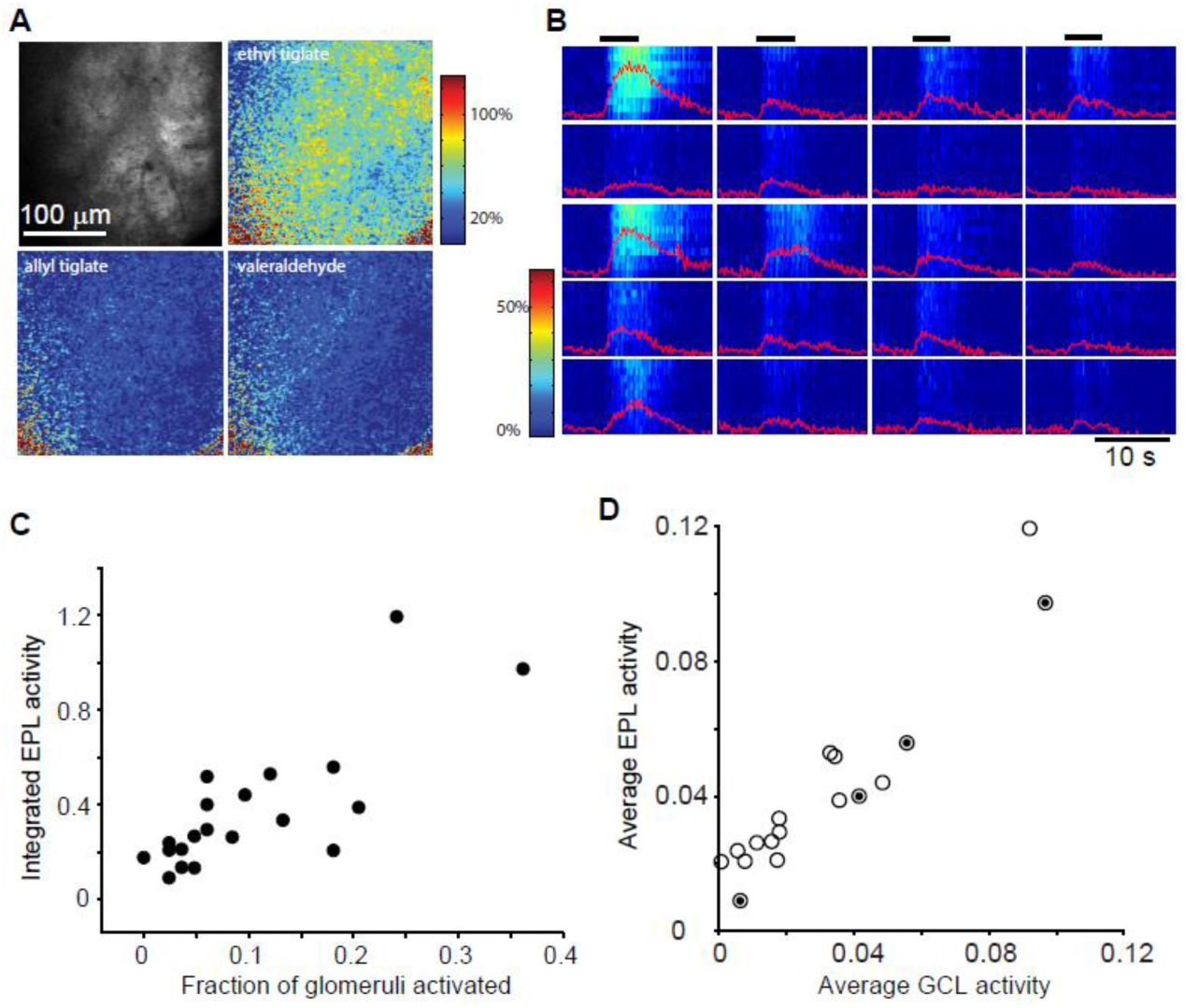
Responses in apical dendrites of GCs. A. Example showing apical dendritic responses with dense labeling of GABAergic neurons with GCaMP3. Individual GC dendrites cannot be discerned, but overall fluorescence changes represent responses from populations of GC dendrites. B. Time course of responses of 10 regions of interest within the imaged field of view for all 20 odors, along with the average traces, are shown. C. Integrated dendritic activity is strongly correlated with density of glomerular activation. D. Average activity in the EPL is also highly correlated with average somatic activity of GCs measured in the same experiment. Circles with dense interior are 4 concentrations of allyl tiglate.

Examining average responses in the EPL is equivalent to recording local field potentials, and the dense labeling of many GCs did not allow us to resolve individual dendrites. To examine responses in individual dendritic segments of single GCs, we turned to sparse labeling using lentiviral infection (Figure 8A). Imaging a region of interest that contained many dendrites, we could readily detect odor-evoked fluorescence increases (Figure 8A,B; Supplementary Movie 4).

**Figure 8:**
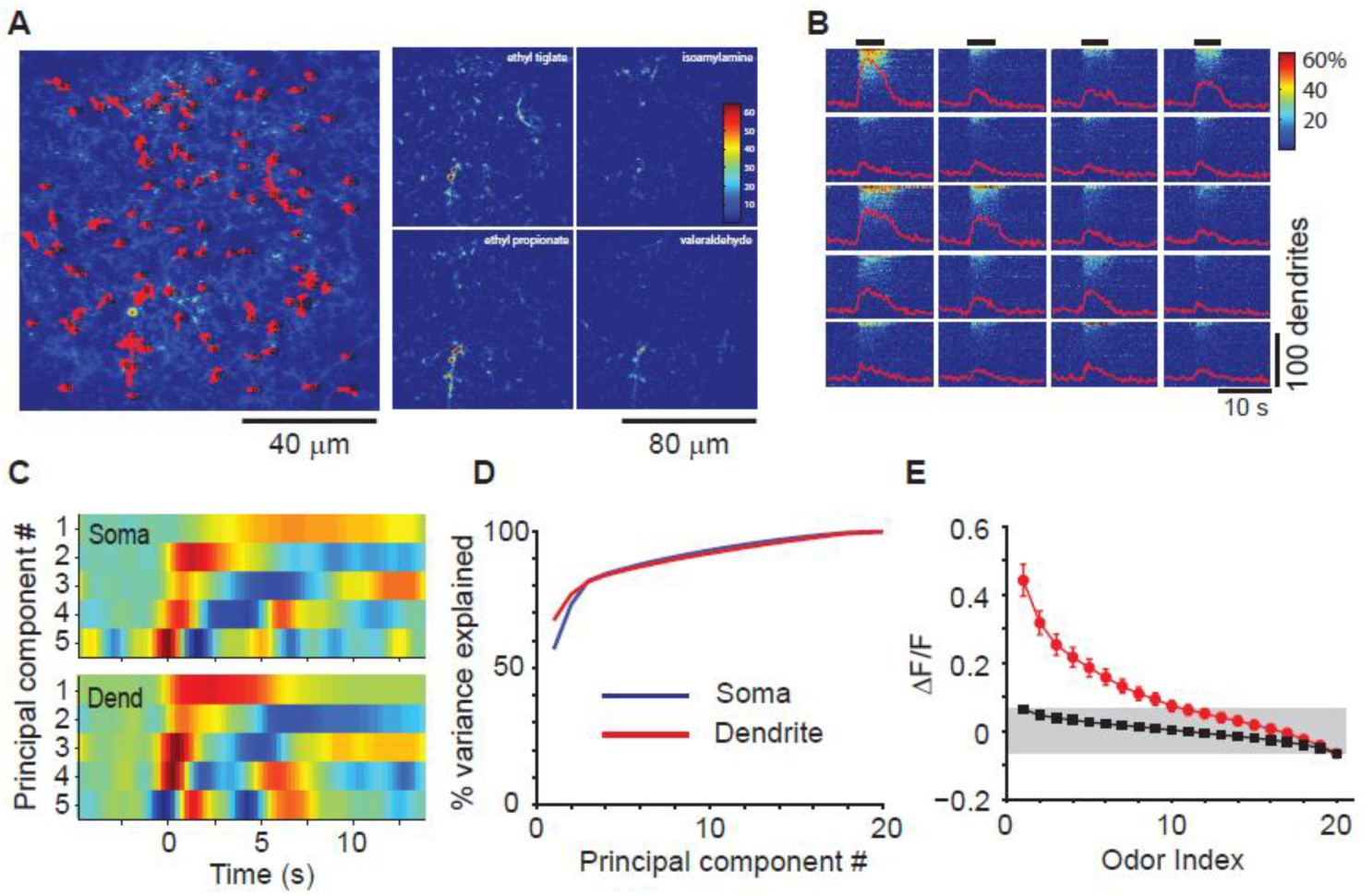
Responses in individual dendritic segments. A. Sparse labeling of GCs using lentiviral injections into the OB allowed individual GC dendrites to be imaged (left) and their responses to odors measured (right). Only a fraction of the regions of interest are highlighted in red in the left image for clarity. B. Time course of responses from 25 randomly chosen dendritic segments to 20 odors, arranged in descending order of response amplitudes for each odor. C. PCA of time course of dendritic responses revealed a faster dynamics of dendritic responses than those for somata. D. Most of the variance in time course of dendritic responses was accounted for by the first few components, as for GC somata (reproduced from Figure 3A). E. The average rank-ordered response amplitude distribution for dendrites revealed that they were generally more responsive than GC somata (compare Figure 4E).

Individual dendritic elements respond to different subsets of odorants used; conversely some odorants evoked sparse responses and others denser activity (Figure 8B). To examine the variation in temporal dynamics of responses, we performed principal component analysis in the same way as we did for somatic responses (Figure 8C). We found that much of the variance of the data could be accounted for by the first few PCs (Figure 8D). Although similar number of PCs were required to explain a given degree of variance for dendritic and somatic responses (Figure 8D), the PCs had systematically faster time courses for dendrites than for GC somata (Figure 8C).

We wondered whether dendrites would be more responsive than somata, since subthreshold synaptic activity could become visible in dendrites through local calcium rises. The number of odors activating a given piece of dendrite was indeed higher for dendrites, compared to somata (Figure 8E; compare with Figure 4E). Rank ordering odor responses for each dendrite and averaging across all dendrites revealed that a dendrite responded to 8 out of 20 odorants on average, compared to 3 out of 20 for somata.

### Local dendritic responses

We next sought to obtain direct evidence for dendritic responses that are independent of somatic responses. To this end, we labeled GCs sparsely using lentivirus injections and imaged the dendrites as well as cell body of visualized GCs (Figure 9A). The soma and dendrites of the same neuron were identified by taking z-stacks of signals in the red channel. Comparing the responses of dendrites and soma revealed that many odors could activate both soma and dendrites (Figure 9B). However, there were also dendritic responses to some odors, with no corresponding activation of somatic responses (*, Figure 9B). This loss of response in the soma was not simply a result of different detection thresholds for dendrites and soma because a given level of dendritic response could either be accompanied by a somatic response (arrow, Figure 9B) or not (*, Figure 9B).

**Figure 9.**
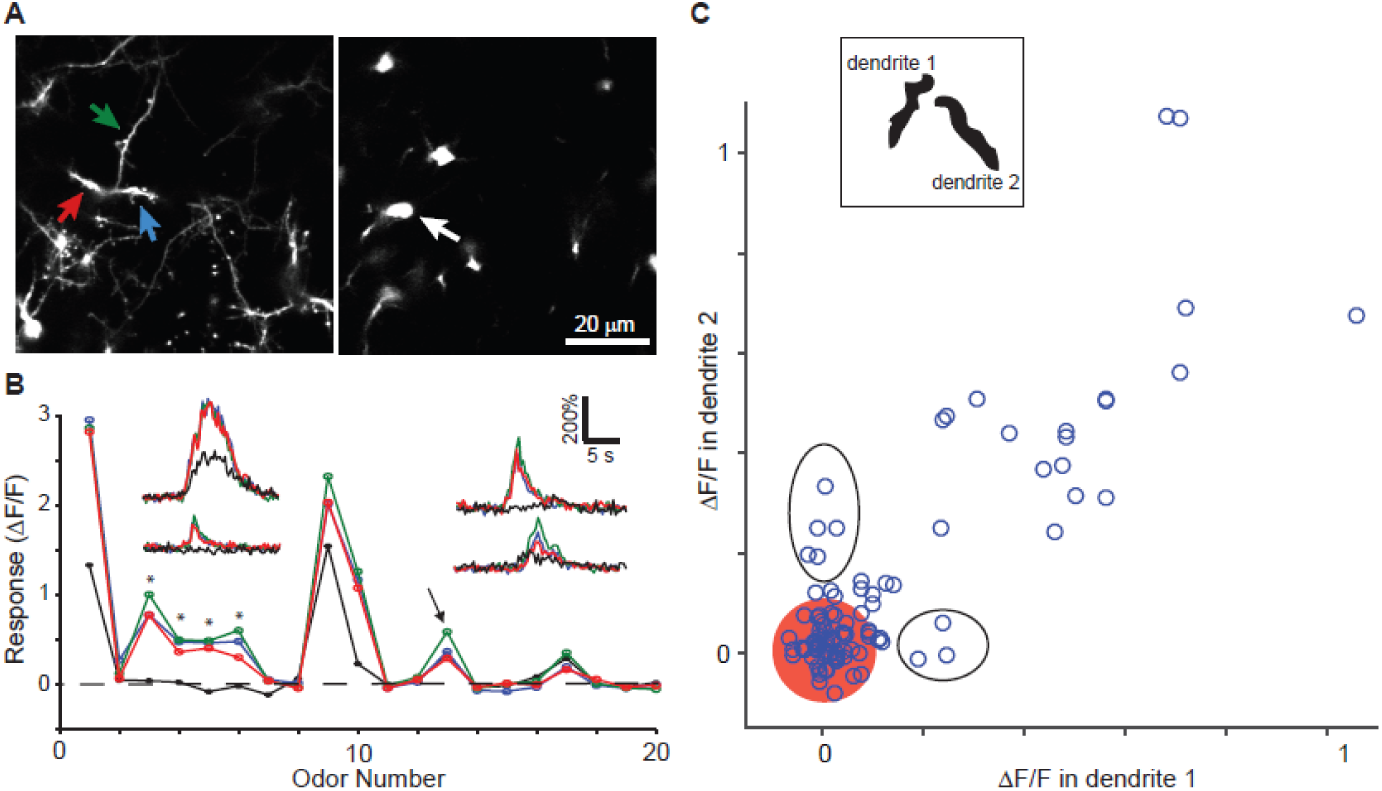
Dendritic responses can be local. A. Dendrites and soma belonging to the same GC were identified through 3-dimensional reconstructions using the dTomato channel. B. Average odor responses for all 20 odors in 3 dendritic segments and the corresponding soma. Dendritic responses to odors 3-6 were not accompanied by somatic responses (asterisks), but a similar magnitude response to odor 13 was accompanied by a somatic response. Insets show time course of odor responses of the 4 regions is shown for 4 selected odors. C. Fluorescence change in pairs of dendrites of the same GC plotted against each other for all 20 odors. Red circle centered at the origin indicates 2 standard deviations of the noise, and ellipses highlight points representing selective dendritic responses that are well above noise.

Our scanning system did not allow us to scan at two different depths simultaneously, forcing us to image somatic and dendritic responses in different trials. Therefore, we focused our efforts on characterizing responses of multiple dendrites of the same GC imaged simultaneously. We looked for odor-evoked responses in an individual dendrite of a GC, in the absence of a response in a different dendrite of the same GC. We identified pairs of dendrites belonging to the same GC and displayed the response of one dendrite against a second dendrite for all 20 odors. Local and selective responses can then be discovered as outliers from the cloud of points along the diagonal (Fig 9C). The experimental yield was lower since sparse labeling was necessary for these experiments (to trace dendrites from an identified GC). We found that 8 out of 34 events were local in the 5 pairs of dendrites we successfully identified that responded to at least one odor (Fig 9C).

These data indicate that local dendritic responses can be occasionally recorded in vivo, demonstrating the possibility of independent dendritic processing.

## Discussion

The function of GCs in the OB has been a matter of speculation for many decades. The biophysical properties of these cells and the reciprocal synapses with M/T cells have been studied in great detail (Egger et al., 2003; Isaacson and Strowbridge, 1998; Jahr and Nicoll, 1980; Rall et al., 1966; Schoppa et al., 1998; Shepherd et al., 2007), but their role in odor processing has been studied far less (Abraham et al., 2010; Fukunaga et al., 2014; Nunes and Kuner, 2015). Here, we have shown that (i) the sparsity of GC population response has a strong relation to the extent of activation of glomerular inputs, (ii) activity of individual GCs can have complex, non-monotonic relationship to stimulus intensity, (iii) functionally similar GCs have some degree of non-random spatial organization of GC responses, (iv) GC dendrites can respond independently to odors, potentially allowing local inhibition. Together, these findings suggest a specific role for GCs, distinct from general-purpose operations such as divisive normalization where participating interneurons indiscriminately integrate activity across many information channels.

### Some caveats and justifications

Before discussing the implications of our study, it is important to acknowledge some caveats and rule out some concerns. First, like most other imaging methods, the signal-to-noise ratio does not approach spike recordings using electrodes, leading to an underestimation of neural activity. Our experiments were done with GCaMP3 (Tian et al., 2009), because this was the best available sensor when the project was conceived and implemented. Although new sensors have better sensitivity (Chen et al., 2013), our main conclusions are still valid since all comparisons of importance were relative, and neither the absolute amplitude of responses nor the absolute number of responsive cells affect our conclusions. In addition, some of the key results were corroborated by experiments using a more sensitive indicator GCaMP5 (Supplementary Figure 3).

A second, related concern is whether the expression level of the indicator biases responses in some way. This concern is significantly alleviated by our observation that there was no correlation between the resting fluorescence of GCs and the maximum response (Supplementary Figure 2), which could have arisen from increased calcium buffering with increased expression levels (Helmchen et al., 1996; Maravall et al., 2000). We also found that a majority of cells in the imaged region are capable of responding to stimuli, as observed during application of bicuculline.

A third caveat is that we did not distinguish among potentially different classes of GCs – for example, in terms of depth of somata (Kelsch et al., 2007; Mori et al., 1983; Orona et al., 1983), neurochemical identity (Davis et al., 1982; Parrish-Aungst et al., 2007) or their age (Kelsch et al., 2007; Lemasson et al., 2005). These will be the focus of future studies.

Fourth, the temporal resolution of signals (both due to the imaging speed and the time constants of the calcium indicator) limited our analysis to time scales of fraction of seconds, rather than milliseconds. Future experiments using speedier and more sensitive probes may allow us to extract faster modulation of activity, and examine its relation to breathing/sniffing.

Finally, there is a question whether imaging in anesthetized animals affects the conclusions made. Two recent studies have indicated that GCs become more responsive to odorants in awake animals (Cazakoff et al., 2014; Kato et al., 2012). We confirmed this finding in our preparation (Supplementary Figure 4). In addition, we found that the increased responsiveness is due to larger responses in individual GCs in awake animals. Our data, combined with previous work, indicates that the overall pattern of activity in awake animals can be approximated by a scaled version of activity in anesthetized animals and therefore does not affect the basic conclusions in our study.

### GC responses are diverse

The ability to record the activity of a large number of GCs allowed us to characterize their response features in a comprehensive manner. Electrophysiological recordings afford excellent signal-to-noise ratios, but have offered limited sample size because of technical challenges of recording from small neurons in vivo. They have, nevertheless, pointed to diverse temporal dynamics (Cang and Isaacson, 2003; Labarrera et al., 2013; Tan et al., 2010; Wellis and Scott, 1990). A previous imaging study also noted diverse temporal dynamics in responses, but focused on response onsets (Kato et al., 2012), which could be related to delayed activation seen in slice experiments (Kapoor and Urban, 2006). Here, we performed principal component analysis of response dynamics and found that the diversity of temporal responses can be accounted for by a small number of primary shapes. In addition to previously reported temporal shapes of transient responses, we found evidence for accumulating or sustained responses (Figure 3B). Temporal coding is thought to be prevalent in the vertebrate OB and its insect analog, arising through bulbar circuit dynamics (Cury and Uchida, 2010; Dhawale et al., 2010; Friedrich and Stopfer, 2001; Laurent, 2002; Raman et al., 2010; Shusterman et al., 2011). Temporally diverse responses in GCs can sculpt mitral cell activity in complex ways, potentially increasing the coding capacity (Giridhar and Urban, 2012; Tripathy et al., 2013).

We also found evidence for inhibitory responses to odorants in GCs. Although many previous electrophysiological studies have not reported inhibitory responses to odorants in anesthetized animals, inhibitory currents could sometimes be recorded under voltage clamp (Cang and Isaacson, 2003; Labarrera et al., 2013). This suggests that inhibitory currents were present, even when the overall responses were dominated by excitation. Inhibitory responses were abolished by blockade of GABA_A_-mediated synaptic transmission. This finding, combined with the observation that inhibition typically appears later than excitation (Figure 3D), indicates that intrabulbar circuit mechanisms underlie inhibitory responses. The source of functional inhibition is likely to be deep short axon cells (Arenkiel et al., 2011; Schneider and Macrides, 1978), but other sources such as GCs themselves could also contribute.

Another novel finding of our study is that responses of GCs are not strictly monotonic as odorant concentration increases. Such non-monotonic responses were not observed in the presence of bicuculline, indicating that intrabulbar inhibition leads to this relation. Responses of principal cells to increasing concentrations have often been presented as being monotonic and saturating (Davison and Katz, 2007; Tan et al., 2010), but many other studies have found complex, non-monotonic dependencies of responses (Meredith, 1986; Wellis et al., 1989). Non-monotonic responses in bulbar neurons (including GCs) may arise because additional, spatially disperse glomeruli will be recruited with increasing odorant concentration (Meister and Bonhoeffer, 2001; Rubin and Katz, 1999), leading to a shift in the balance of excitation and inhibition impinging on a given cell. For this reason, increasing concentration should not be seen as analogous to increasing stimulus intensities in other modalities, where the stimulus locus is held fixed.

### GC response density scales with input

Another key finding of our study is that the response of GCs as a population scales well with the density of glomerular input. Electrophysiological recordings have hinted at weak responses to odorants in anesthetized rodent (Cang and Isaacson, 2003; Cazakoff et al., 2014; Wellis and Scott, 1990), but the sparseness of responses to a variety of odorants has not been studied systematically (but see (Tan et al., 2010)). Recent imaging studies have also noted that the response of GCs are very sparse in anesthetized animals, again without characterizing the glomerular activation patterns of these odorants (Kato et al., 2012). However, any discussion of the sparseness of responses of a population of neurons has to take into account the stimulus space. For example, in the visual system natural scenes lead to sparser encoding by cortical neurons than high contrast bars or gratings (Froudarakis et al., 2014; Vinje and Gallant, 2000; Weliky et al., 2003).

In this study, we examined the responses of GCs in relation to the inputs to the OB. We find that even in the anesthetized animal, the density GC responses as a population changes gradually with increasing number of activated glomeruli. Graded increase in the number of GCs could offer increasing inhibition to MCs as inputs become stronger, reducing saturation. It is, therefore, important to indicate how densely any particular odor stimuli activate glomeruli when quantifying response sparseness. In addition, our results also indicate that GCs are capable of responding to sensory inputs even when there is likely to be minimal feedback from higher brain areas, which is thought to be more prominent in the awake state (Boyd et al., 2012, 2015; Otazu et al., 2015; Restrepo et al., 2009).

Odor-evoked responses in GCs become denser in the awake compared to anesthetized state (Cazakoff et al., 2014; Kato et al., 2012). The reasons for this differential response sensitivity of GCs include state-dependent neuromodulation (Kiselycznyk et al., 2006; Ma and Luo, 2012; Petzold et al., 2009; Rothermel et al., 2014), differing extent of functional cortical feedback (Boyd et al., 2012, 2015; Markopoulos et al., 2012; Otazu et al., 2015) and the presence of tonic inhibition (Labarrera et al., 2013). We were able to confirm in our experiments that GCs are indeed more responsive to odor stimulation in the awake compared to anesthetized state (Supplementary Figure 4). However, the basic conclusions of our studies in anesthetized animals are not affected since the relation between input structure and response density is preserved, and we found non-monotonic responses of some GCs to increasing odor concentration (Supplementary Figure 4). In addition, we found that responses in awake animals are variable because of fluctuations in stimulus sampling due to sniff modulation, which made it harder to characterize response features from limited number of trials. We feel that responses in anesthetized animals are governed more by feedforward bulbar circuitry, establishing a ground state for further analysis in more awake and attentive states, where more complex higher order feedback will lead to variable effects. Independent of the variation in the degree of sparseness in different brain states, our findings demonstrate that firing sparseness is related to the statistics of the sensory input itself.

### Non-random spatial pattern of activation of GCs

Mitral cells with their primary dendrites in the same glomerulus are anatomically clustered in a small region, simply by geometric constraints (Buonviso et al., 1991). Their lateral dendrites, however, span hundreds of microns and therefore become enmeshed with dendrites of mitral cells belonging to other glomeruli. Therefore, GCs activated by sister mitral cells can potentially be spatially distributed and intermingled with GCs activated by other dissimilar mitral cells (Murthy, 2011). Despite this distributed organization, there is anatomical evidence for potential clustering of GCs, but this is based on poorly understood trans-synaptic viral transport (Willhite et al., 2006). Experiments using c-fos labeling to quantify neural activation have also suggested clustered activation of GCs (Guthrie et al., 1993), but this method has unknown sensitivity. Using a real-time imaging method that has much better sensitivity and resolution, we found evidence for local functional clustering of GCs.

Initial analysis of the spatial pattern of activated GCs across all odorants revealed little order. However, any local order related to glomerular identity may be obscured because a single odorant can activate spatially distributed glomeruli and neighboring glomeruli can be functionally distinct (Bozza et al., 2004; Ma et al., 2012; Soucy et al., 2009). We reasoned that odorants that activated glomeruli sparsely (ideally a single glomerulus) could reveal underlying spatial order of activated GCs. In line with this reasoning, we found that pairwise separation of active GCs was smaller for odors that that activated glomeruli very sparsely, than for odors that had dense activation. This provides strong evidence that when a single glomerulus is stimulated, GCs that are activated are clustered spatially.

Does this clustering imply that there is selective synaptic connectivity, such that GCs in a small region get higher than chance inputs from sister mitral cells? Although we cannot rule out such selective functional synaptic connectivity, a simpler situation based on purely geometric considerations could allow clustered GC activation (Murthy, 2011). This is because, as sister mitral cells diverge, the chance that multiple lateral dendrites of sister MCs will activate neighboring GCs decreases as a square of the distance. Therefore, even with uniform connection probability along the lateral dendrite, a clustering of activated GCs will be observed. Future experiments combining functional and anatomical characterization will be necessary to disambiguate these possibilities.

It is not clear what such a clustering of GCs may accomplish. One possibility is that a cluster of functionally similar GCs can target the proximal dendrites of mitral/tufted cells within a local region to create a coarse and dynamic functional cell assembly (Yu et al., 2013).

### Local dendritic processing

It has long been realized that the reciprocal synapses between M/T cells and GCs offer substrates for localized function (Egger et al., 2005; Shepherd et al., 2004; Yu et al., 2013). Unlike axon-dendritic synapses, which generally rely on action potential propagation, local dendrodendritic synapses in the OB and other brain circuits such as the retina, can react to local depolarizations (Euler et al., 2002). In vitro slice work has provided strong evidence the dendrites of GCs can sustain local depolarization as well as global activity, presumably due to action potentials invading the entire dendritic tree (Egger et al., 2003, 2005). Clear evidence for local dendritic calcium events and accompanying dendritic spike events recorded in the soma was also presented in the more intact frog nose-brain preparation (Zelles et al., 2006). Whether such local depolarizations occur *in vivo* in response to odor stimuli has remained unknown. Here, we offer strong evidence for the existence of local depolarization in GC dendrites *in vivo.*

First, we found that the average odor tuning curves for dendrites were broader than those for GC somata. This difference is unlikely to be simply due to differences in calcium buffering, since a careful study in slices has revealed no significant heterogeneity of calcium handling in GCs (Egger and Stroh, 2009). Second, the temporal dynamics of responses in dendrites were, at a population level, different from those in soma. Third and most directly, some odors could evoke calcium rises in dendrites without any detectable responses in the corresponding GC soma. This was not simply due to reduced sensitivity in the soma because other odors triggering fluorescence increases of similar magnitude in the dendrites can lead to detectable somatic signals as well.

What function could local dendritic activity in GCs have? Since many of the dendrodendritic synapses are reciprocal (Shepherd et al., 2004), local depolarization will mainly affect only those presynaptic mitral cells that caused the GC depolarization, leading to reciprocal or autoinhibition. By contrast, global depolarization of a GC can lead to lateral inhibition that affects other mitral/tufted cells.

### GCs provide specific inhibition

It is now clear that interneurons in many regions of the brain are diverse (Ascoli et al., 2008; Fishell and Rudy, 2011), and each class of interneurons may have specific functional role (Fishell and Rudy, 2011; Jadzinsky and Baccus, 2013). In the mouse retina, for example, there are more than 20 types of amacrine cells and their roles are beginning to be dissected (Jadzinsky and Baccus, 2013). In the OB, there is increasing evidence that some interneurons, including certain classes of glomerular-layer interneurons and parvalbumin-positive neurons in the external plexiform layer – may play a role in gain control or normalization (Kato et al., 2013; Miyamichi et al., 2013). Normalizing function has been well-studied in many regions of the brain and has attractive theoretical and conceptual implications (Carandini and Heeger, 2012; Isaacson and Scanziani, 2011). However, for many circuit computations, more specific inhibition is necessary. For example, in the retina, the classical lateral inhibition gives rise to edge detection (Gollisch and Meister, 2010; Jadzinsky and Baccus, 2013).

GCs in the OB have long been postulated to offer specific inhibition (Cleland, 2010; Koulakov and Rinberg, 2011; Shepherd et al., 2007; Wick et al., 2010) and our study presents several experimental observations supporting this postulate. First, the existence of inhibitory responses in GCs is not readily predicted by models of global normalization. Second, non-monotonic responses in GCs with increasing concentration support a model of OB circuit function where there are competitive interactions among these cells, leading to dynamic and flexible inhibition of principal cells (Cleland, 2014; Koulakov and Rinberg, 2011). Again, this is in contrast to interneurons that function to integrate increasing inputs in a monotonic manner, and provide normalizing inhibition to principal cells (Kato et al., 2013; Miyamichi et al., 2013). Finally, the existence of local dendritic responses independent of the GC soma also strongly suggests that GCs can influence their targets more locally. Collectively, our data adds support to the idea that GCs mediate local and feature-specific inhibition. Such local dendritic responses in amacrine cells have been shown to be a key feature of direction selective responses in the retina (Euler et al., 2002; Jadzinsky and Baccus, 2013). In the OB, our data suggest that GCs do not simply integrate the activity of all glomerular channels, but can selectively pool information from specific glomeruli. A careful consideration of odorant stimulus features and the glomeruli they activate may catalyze experiments illuminating GC function in the OB.

## Materials and Methods

### Animals and Surgery

Adult, 8 to 12 week-old male C57BL/6 (Charles River), VGAT-ires-Cre (Jackson Laboratory strain *Slc32a1*^*tm2(cre)Lowl*^/J) or OMP-GCaMP3 mice (Isogai et al) were anesthetized with an intraperitoneal injection of ketamine (90 mg/kg) and xylazine (10 mg/kg) and body temperature was maintained at 37°C by a heating blanket (Harvard Apparatus). For viral injections a small craniotomy was made over the olfactory bulb and 100 μl of AAV2/9 or 300 μl of lentivirus was injected at a depth of 700 μm below the dural surface through a glass micropipette attached to a nanoinjector (MO-10, Narishige). To create an optical window an aluminum plate was glued to the skull, and the olfactory bulbs were exposed by making a craniotomy on either hemisphere using a dental drill. The surface was kept moist with artificial cerebrospinal fluid (aCSF; 125 mM NaCl, 5 mM KCl, 10 mM Glucose,10 mM HEPES, 2 mM CaCl2 and 2mM MgSO4 [pH 7.4]). Before imaging, 1 % agarose in aCSF was placed on the bulb and the window was closed with a coverslip. To block inhibition a small cut through the dura was made at the periphery of the craniotomy and a solution of 1% agarose in aCSF containing 100 μM (-)-Bicuculline methiodide (Tocris) was applied to the bulbar surface. All experiments were approved by the Standing Committee on the Use of Animals in Research of Harvard University.

### Viral vectors

Granule cells were labeled with AAV2/9 expressing tdTomato and GCaMP3 under control of the CMV early enhancer/chicken β actin (CAG) promoter. Bicistronic expression was achieved by using a ribosomal skip site (T2A) between the two fluorophores. For conditional expression in VGAT-Cre mice we used AAV carrying floxed GCaMP3. AAV vectors were produced by the University of Pennsylvania Viral Vector Core. For sparse labeling we made use of a Tet-Off lentiviral system (Hioki et al., 2009) with one construct expressing the transactivator tTAad under control of a synapsin promotor and a second construct expressing tdTomato-T2A-GCaMP3 under control of a TRE promoter. VSV-G pseudotyped lentiviral vectors were produced by transfection of human embryonic kidney cells (HEK293) with third-generation lentivirus plasmids using lipofection (Mirus). Supernatant was collected 48 h after transfection and concentrated using centrifugal filters (Millipore).

### Immunohistochemistry

Three weeks after surgery, virus-injected mice were deeply anesthetized with a Ketamine (180 mg/kg i. p.)/Xylazine (20 mg/kg, i.p.) mixture and perfused transcardially with 20 ml of PBS (pH 7.4) first, followed by 50 ml of 4% paraformaldehyde in 0.1 M phosphate buffer (pH 7.4). Brains were removed and cut into 70 μm-thick sagittal sections with a vibratome (Leica). Sections were permeabilized and blocked with a solution containing 0.1% Triton X-100 (Tx, Fisher), 0.5% carrageenan (Sigma), and 2.5% goat serum in PBS for 1 h, and incubated overnight with primary antibody against GFP conjugated to Alexa 488 dye (Invitrogen) diluted 1:1000 in blocking solution. Sections were imaged with a confocal microscope (LSM 710, Zeiss).

### In vivo imaging

A custom-built two-photon microscope (Wienisch et al., 2012) was used for *in vivo* imaging. Fluorophores were excited and imaged with a water immersion objective (20×, 0.95 NA, Olympus) at 950 nm using a Ti:Sapphire laser (Mai Tai HP, Spectra-Physics). Frame rates were typically 4 Hz. Image acquisition and scanning were controlled by custom-written software in Labview. Emitted light was routed through two dichroic mirrors (680dcxr, Chroma and FF555-Di02, Semrock) and collected by 2 photomultiplier tubes (R3896, Hamamatsu) using filters in the 500-550 nm range (green channel, FF01-525/50, Semrock) and 572-642 nm range (red channel, FF01-607/70, Semrock).

To identify sparsely labeled granule cells that allow imaging of both their soma and dendrites the laser was tuned to 1040 nm and the red channel was used to record a detailed z-stack with 1 μm step size. The entire dendritic tree of a given GC was then examined to ensure that it could be clearly separated from processes of neighboring cells and faithfully traced to its corresponding soma. After choosing one imaging plane containing the soma and one imaging plane containing several branches of the same dendrite the laser was tuned back to 950 nm for imaging odor-evoked activity in either plane.

### Odor stimulation

Following monomolecular odorants (Sigma) were used as stimuli and delivered by a custom-built 20 channel olfactometer controlled by custom-written software in Labview (National Instruments): ethyl tiglate (1), ethyl acrylate (2), valeric acid (3), allyl butyrate (4), isoamylamine (5), 2-methoxypyrazine (6), pyrrolidine (7), piperidine (8), allyl tiglate (9), valeraldehyde (10), isoamyl acetate (11), heptanal (12), iseugenol (13), ethyl propionate (14), 1-pentanol (15), 2-heptanone (16), p-anis aldehyde (17). Individual odors were presented for 5s with an interstimulus interval of 60 s. Odorants were used at a nominal volumetric concentration of 1 % (v/v) in mineral oil. For experiments testing the concentration dependence of responses, allyl tiglate was used at the following nomimal dilutions (v/v): 0.1 %, 0.5 %, 1 % and 10 %. The actual concentration ranges at the nose of the animal was estimated using measurements from a miniPID detector (Aurora). These values were then expressed as log dilution.

### Data analysis

Data were analyzed offline using custom-written scripts in MATLAB (Mathworks). We selected regions of interest for analysis from averages of 100 or more individual images (to increase signal to noise ratio). Somata or dendritic segments were chosen based on fluorescence in the red (dTomato) channel. The average intensity in the green (GCaMP) channel was calculated for each ROI, for each frame and for each odor, leading to a 3D matrix of numbers.

A response value for each cell-odor pair was calculated as the average ΔF/F value over the 5 seconds of odor presentation. We performed receiver operating characteristic (ROC) analysis on our data and found a ΔF/F response threshold of 0.05. Since the noise distribution and signal distribution are highly overlapping (Figure 2c), this threshold results in underestimating the number of responding cells. However, since most of the analysis relies on relative numbers, the exact number of responding cells will not affect our conclusions.

### Temporal dynamics

Principal component analysis of the time course of responses was performed in Matlab (Mathworks) using centered data and singular value decomposition. The delay in the onset of responses was calculated from reconstruction of the time course using the first 5 principal components (which smoothed the data). Onset time was defined as the first time the ΔF/F value deviated from the baseline, if at least 3 subsequent points were 3 standard deviations above baseline fluctuation.

### Rank ordered odor tuning

To facilitate averaging of odor tuning curves across all cells, we ordered individual tuning curves in descending order of amplitudes and averaged them across all cells. This average ordered tuning curve needs to be compared with a control tuning curve to estimate the number of odors an average GC responds to. The control curve was estimated as follows. We obtained a sample of 20 entries from a Gaussian noise distribution (Figure 2c) and rank ordered them as if they were responses. We repeated this procedure 300 times to mimic many GCs, and averaged the rank-ordered tuning curves (obtained from a noise distribution). The maximum and minimum values of this null tuning curve gave us noise bounds, and the number of entries in the actual response tuning curve that exceeded these bounds gave us an estimate of the average number of odors that activated a GC.

### Sparseness measures

To quantify the sparseness (or density) of activity of GCs, we estimated two measures: lifetime sparseness and population sparseness. Lifetime Sparseness quantifies the extent to which a given neuron is activated by different odor stimuli (Willmore and Tolhurst, 2001). If all stimuli activate the observed element rather uniformly, the lifetime sparseness measure will be close to 1, and if only a small fraction of the stimuli activate the unit significantly, this metric will be close to 0.

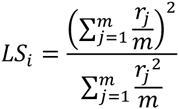

Where:

*m* = *number of odors*, *r*_*j*_ = *response of neuron A to odor j*

*i* refers to the index of the glomerulus A or cell body A, for which lifetime sparseness is calculated. Population Sparseness quantifies the fraction of elements (glomeruli or mitral cells) in the imaged field of view that respond to a given odor from the stimulus panel presented (Willmore and Tolhurst, 2001).

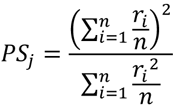

Where: *n* = *number of neurons*, *r*_*i*_ *= response of neuron A to odor j*

*j* refers to the index of the odor stimulus in the panel for which the population sparseness is calculated.

### Response similarity and spatial separation

To quantify the similarity of response between pairs of GCs (indexed as A and B), we used the centered (Pearson) correlation coefficient:

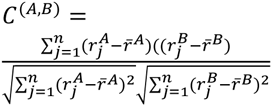

Where

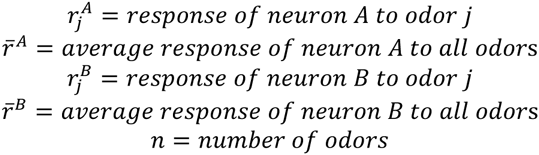

We plotted the relationship between similarity and physical separation and averaged the similarity values by binning the data. Expected values for a random distribution were calculated by assigning randomly shuffled tuning curves to different GCs, and calculating pair-wise similarities.

### Spatial distribution of responding GCs

The pair-wise separation of active GCs was calculated for each odorant, and the average separation for each odor was plotted against the extent of glomerular activation (Figure 6D). The expected separation was estimated by sampling random pairs of GCs (from all the responding GCs within the imaged field of view). The effect of small sample numbers in potentially biasing the relation between glomerular sparseness and pairwise GC separation was assessed in the following way. We sampled randomly small number of GCs (equivalent to the numbers experimentally found for sparse odors) from the entire population of active GCs in a given experiment, and calculated the average pairwise separation. We repeated this sampling procedure multiple times to examine the scatter in the estimate of pairwise separation from small numbers of GC pairs. We found that the distribution of this average had high variance, but the average separation was close to the value predicted for all GCs (127 μm) (Supplementary Figure 7).

## Acknowledgment

We thank the members of our group for fruitful discussions and comments on the manuscript. This research was supported by a HFSP Long-term fellowship to MW and funds from Harvard University and the NIH.

## Contributions

MW and VNM designed the study, MW performed all experiments, MW and VNM analyzed the data and wrote the paper.

## Conflict of Interest

The authors declare no competing financial interests.

